# The Orbitofrontal Cortex in Temporal Cognition

**DOI:** 10.1101/2020.10.16.343178

**Authors:** Juan Luis Romero Sosa, Dean Buonomano, Alicia Izquierdo

## Abstract

One of the most important factors in decision making is estimating the value of available options. Subregions of the prefrontal cortex, including the orbitofrontal cortex (OFC), have been deemed essential for this process. Value computations require a complex integration across numerous dimensions, including, reward magnitude, effort, internal state, and time. The importance of the temporal dimension is well-illustrated by temporal discounting tasks, in which subjects select between smaller-sooner versus larger-later rewards. The specific role of OFC in telling time and integrating temporal information into decision making remains unclear. Based on the current literature, in this review we reevaluate current theories of OFC function, accounting for the influence of time. Incorporating temporal information into value estimation and decision making requires distinct, yet interrelated, forms of temporal information including the ability to tell time, represent time, create temporal expectations, and the ability to use this information for optimal decision making in a wide range of tasks, including temporal discounting and wagering. We use the term ‘temporal cognition’ to refer to the integrated use of these different aspects of temporal information. We suggest that the OFC may be a critical site for the integration of reward magnitude and delay, and thus important for temporal cognition.

## Role of OFC in Learning and Decision making

The Orbitofrontal Cortex (OFC) has long been associated with the updating of stimulus–reward associations (Jones & Mishkin, 1972; Klein-Flügge, Barron, Brodersen, Dolan, & Behrens, 2013; Ostlund & Balleine, 2007) and with encoding the current value of rewards (Gottfried, Doherty, & Dolan, 2003; Roesch, Taylor, & Schoenbaum, 2006; Tremblay & Schultz, 1999). Historically, one of the earliest employed tasks that demonstrated the involvement of OFC in flexible learning and decision making was the stimulus-reward reversal task (Mishkin, 1964). In this task, monkeys were trained to associate a specific stimulus with a reward, over another stimulus that yielded no reward. After monkeys learned these associations, reinforcement contingencies were reversed such that selecting the previously unrewarded stimulus would now yield reward and vice versa. Monkeys with an intact OFC were able to quickly adapt to the new reinforcement contingencies, whereas monkeys with aspiration lesions of OFC were impaired. More recent investigations have resulted in the determination that primate posterior-lateral OFC (and its widespread cortico-cortical connections with anterior insula, and lateral OFC) is a critical substrate of reversal learning (Peter H. Rudebeck, Saunders, Prescott, Chau, & Murray, 2013; Sallet et al., 2020). Indeed there is now also evidence of functional heterogeneity in rodent OFC in supporting reversal learning (Hervig et al., 2019; Alicia Izquierdo, 2017; Verharen, den Ouden, Adan, & Vanderschuren, 2020), and value updating (Bradfield, Dezfouli, van Holstein, Chieng, & Balleine, 2015; Gourley, Zimmermann, Allen, & Taylor, 2016; Malvaez, Shieh, Murphy, Greenfield, & Wassum, 2019).

Another early seminal experiment introduced a phenomenon closely related to reversal learning: “perseveration,” or a failure to disengage from responding to previously rewarded stimuli (Rosenkilde, Rosvold, & Mishkin, 1981). Perseveration has been a focal topic of many psychiatric and translational studies because several human clinical disorders (substance use disorder, anxiety disorders, obsessive-compulsive disorder, etc.) are characterized by “sticky” stimulus-reward associations (Ersche, Roiser, Robbins, & Sahakian, 2008; Remijnse et al., 2006; Ruscio, Seitchik, Gentes, Jones, & Hallion, 2011). An important factor related to both reversal learning and a perseverative phenotype is the causality assigned to a stimulus as predictive of the reward, and indeed there is evidence that lateral OFC plays a crucial role in such credit assignment (Akaishi, Kolling, Brown, & Rushworth, 2016; Noonan et al., 2010; Alexandra Stolyarova, 2018; Walton, Behrens, Noonan, & Rushworth, 2011). OFC supports flexible solving of the structural credit assignment problem, i.e., in determining *which* stimuli reliably predict reward (Akaishi et al., 2016; Noonan et al., 2010). Yet it is also essential to consider how OFC may be involved in learning conditions where there is a delay between the stimulus and the outcome, referred to as the temporal credit assignment problem (Walsh & Anderson, 2011). Recently it has been proposed that OFC may contribute to establishing causal relationships by signaling desirability of the outcome (Grossberg, 2018; P. H. Rudebeck, Saunders, Lundgren, & Murray, 2017) likely via its connections to areas important in incentive value in both rodent and primate species, such as the basolateral amygdala (Gallagher et al. 1999; Baxter et al. 2000; Schoenbaum et al. 2003), ventral striatum (McDannald et al., 2012), and hypothalamus (Petrovich & Gallagher, 2007). Thus, maintaining a causal stimulus-outcome relationship is necessary when a delay between these two events is introduced. Still, it is not well-understood precisely how OFC, or prefrontal cortex generally, supports this ability. Eligibility traces have been proposed as ways to encode delays in reinforcement learning (Sutton & Barto, 1998) and are thought to decay exponentially. OFC could contribute this eligibility trace, although as we note later detail, most discounting models rely on hyperbolic, not exponential, functions in their descriptions. Below we will discuss the possibility that OFC is critical in providing a combined statistic that incorporates both value and time into learning and decision making. As such, it is predicted to be critical in solving the temporal credit assignment problem as well.

Other studies that demonstrate the involvement of OFC in value-based decision making employ reinforcer devaluation procedures (A. Izquierdo, Suda, & Murray, 2004; Pickens et al., 2003; West, DesJardin, Gale, & Malkova, 2011). Such methods usually involve degrading the value of a specific reward by overfeeding the animal to satiety with it (or pairing the reward with an aversive outcome, like illness), and then testing if the subject can update the value of the cue(s) or stimuli associated with that reward. In these experiments, OFC (and amygdala) must be online to encode the new value via experience of the overfeeding, and respond appropriately during testing (West et al., 2011; Zeeb & Winstanley, 2013), c.f. (Fisher, Pajser, & Pickens, 2020). Relevant to our discussion here, for devaluation of a specific reward to occur via satiety, the integration of a largely disproportionate reward magnitude over time is needed.

Another prominent observation is the recruitment of OFC in economic decisions, e.g., value computation depends on other options available for choice, which is correlated with OFC signaling. Early single-unit recording experiments conducted in primate OFC resulted in the demonstration that a subset of neurons fired selectively to different rewards (Rosenkilde, Bauer, & Fuster, 1981), as well as their relative value (i.e., preference), associated with their sensory specificity or unique identities (Thorpe, Rolls, & Maddison, 1983; Tremblay & Schultz, 1999). Padoa-Schioppa and Assad (2006) later showed that the firing rate of OFC neurons corresponds to the economic value of offered and chosen rewards in monkeys, with a similar relationship recently reported in mouse OFC (Kuwabara, Kang, Holy, & Padoa-Schioppa, 2020). Importantly, unlike primates (Rushworth, Noonan, Boorman, Walton, & Behrens, 2011), in rats a choice between offers does not rely on either medial or lateral OFC (M. P. Gardner et al., 2018; M. P. H. Gardner, Conroy, Shaham, Styer, & Schoenbaum, 2017) unless there is a need for a value update or new information about outcome value, assessed by pre-feeding rats to devalue the outcome (M. P. H. Gardner, Conroy, Sanchez, Zhou, & Schoenbaum, 2019).

Collectively, these lines of research demonstrate the importance of OFC in encoding and updating reward but do not explicitly address the contribution of time to decision making. Next, we review the importance of the temporal dimension in decision making and examine the role of OFC in integrating temporal information into value-based decisions.

## Importance of Timing in Naturalistic Decision Making

A fundamental dimension of decision making pertains to time. Humans and other animals prefer earlier rewards over later rewards of the same magnitude, and often smaller-immediate rewards over larger-later rewards. This phenomenon is known as temporal or delay discounting. In the human literature, temporal discounting is often exemplified by the observation that people tend to choose an immediate reward of $100 over a reward of $120 in one month. Interestingly, however, when the same values are placed in the distant future, there is a shift towards more patient preferences: more people choose a $120 option in 13 months versus $100 in 12 months (Frederick, Loewenstein, & O’Donoghue, 2002). These time-dependent decisions are presumably rooted in millions of years of evolutionary pressures for animals to adapt to unique foraging and ecological niches (Murray, Wise, & Rhodes, 2011). Time is a unique resource to all animals as optimal behavior is guided by species- and niche-specific temporal contingencies (Stevens, Hallinan, & Hauser, 2005; Stevens, Rosati, Ross, & Hauser, 2005). For example, brief bouts of foraging for small rewards may decrease the time an animal is exposed to predators, and the amount of energy spent. Similarly, predators must balance the time invested in waiting for prey in a given location and when to “cut its losses.” These processes have been studied by many groups as the explore-exploit tradeoff, for example, in the context of balancing the maximization of information (i.e., knowing where the reward patches are; explore) versus maximizing reward at known patches (i.e., exploit) (Addicott, Pearson, Sweitzer, Barack, & Platt, 2017; Costa & Averbeck, 2020; Gonzalez & Dutt, 2011).

The importance of time as both a resource and a modulator of value implies that decision making must take place in the context of numerous time-dependent factors, including the overall temporal structure of expected reward delays, reinforcement history, replenishment time, and urgency (e.g., current internal states such as hunger or fear of predators). Thus, the brain must first be able to tell time across multiple time scales, store this information, and then integrate it into value computations and decision making. Together we will refer to this array of time-dependent computations as *temporal cognition*. And we propose that circuits involved in decision making, including various areas of the prefrontal cortex, including the OFC, must have access to timing information and perform computations based on the interaction of nontemporal and temporal dimensions, such as the ratio of reward magnitude over delay-to-reward.

## Keeping Time vs. Using Temporal Information

The importance of time to sensory processing, animal behavior, and decision making involve many distinct but interrelated temporal computations, including, telling time (i.e., measuring elapsed time), generating well-timed motor responses, and creating temporal expectations, as well as learning, representing, and storing temporal relationships (Paton & Buonomano, 2018). Together all these components are important for cognitive tasks such as temporal discounting or deciding to stay in, or leave, a reward patch (temporal wagering).

At the first level, temporal cognition requires that the brain have mechanisms to tell time, that is, “clocks” or “timers” that allow animals to distinguish between short and long reward delays, anticipate the arrival of a reward, and decide when to abandon a reward patch. The ability to tell time, and the underlying neural underpinnings, are often studied in the context of explicit and implicit timing tasks (Ameqrane, Pouget, Wattiez, Carpenter, & Missal, 2014; J. T. Coull & Nobre, 2008; A. C. Nobre, Correa, & Coull, 2007). Explicit timing (**Figure 1A)** refers to tasks in which timing is explicitly required for completion of a task, such as discriminating auditory tones of an 800 ms versus a 1000 ms stimulus, generating differentially delayed motor responses in response to two different sensory cues, or producing intricate temporal motor patterns (Slayton, Romero-Sosa, Shore, Buonomano, & Viskontas, 2020; Wang, Narain, Hosseini, & Jazayeri, 2018; Wright, Buonomano, Mahncke, & Merzenich, 1997). Implicit timing tasks (**Figure 1B**) refer to those for which in principle it is not necessary to track time to perform the task but rather learning the temporal structure of the task can improve performance (J. T. Coull & Nobre, 2008; Anna C. Nobre & van Ede, 2018). For example, in a simple foreperiod reaction time task, humans learn the interval between the “ready” and “go” cues, even though they simply need to respond to the go cue (Niemi & Näätänen, 1981). Indeed, humans and animals create a temporal expectation based on the history of the task structure that is built-up across trials, and when target stimuli appear at the expected time, performance is improved and reaction time decreased (Cravo, Rohenkohl, Santos, & Nobre, 2017; Janssen & Shadlen, 2005; van Ede, Niklaus, & Nobre, 2017).

**Figure 1.**
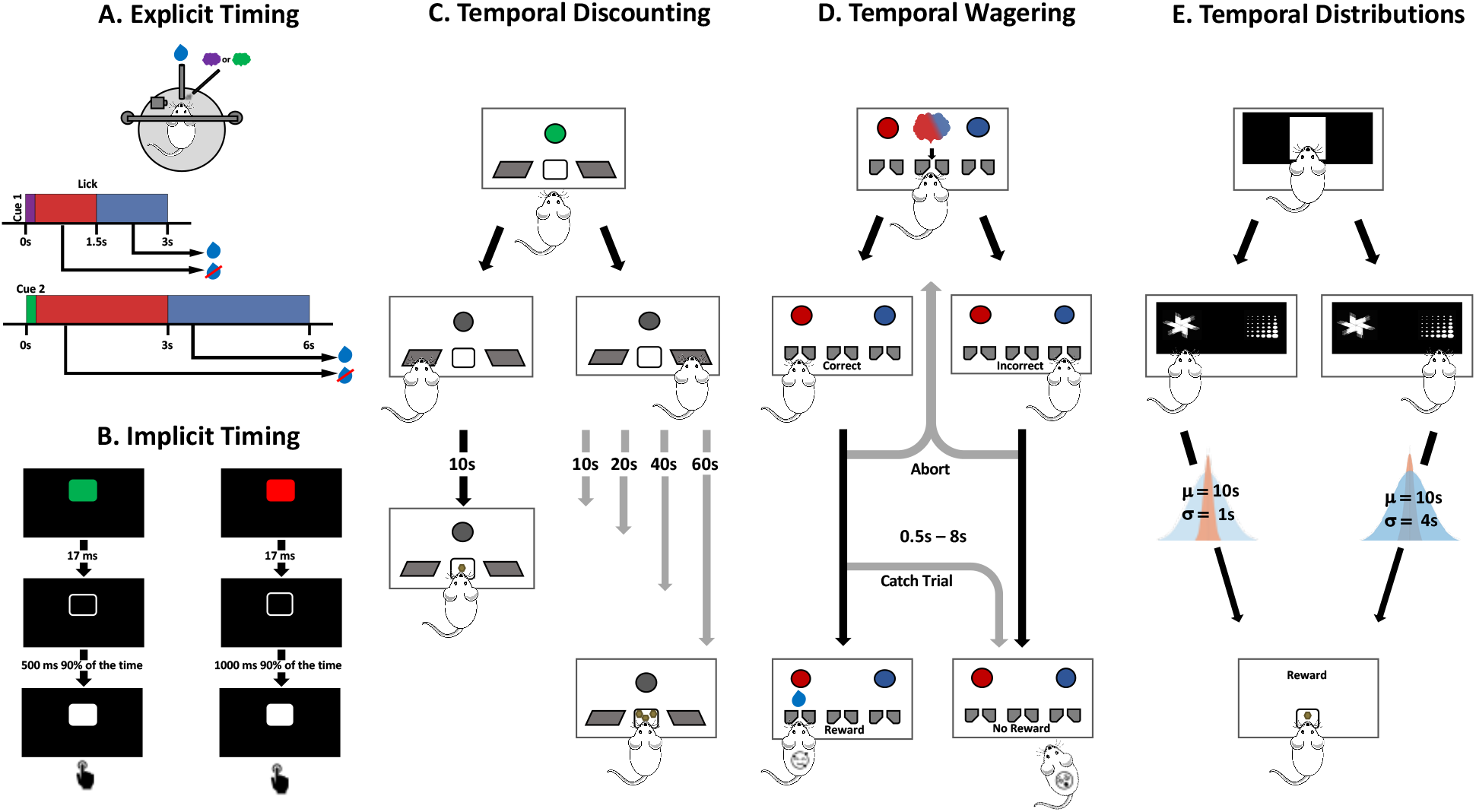
Examples of timing and temporal cognition tasks. **A.** In an explicit timing task, a mouse learns the reward delay associated with each cue and produces anticipatory licking during the appropriate cue-specific interval. **B.** In an implicit timing task each trial may be initiated by a cue (green or red), and humans are asked to respond to a white square. In valid trials each cue is associated with a short or long delay, but in a small number of invalid trials the relationship is reversed. Reaction times are faster in the valid trials even though the task simply requires responding to the target. **C.** Temporal discounting tasks require animals to select between a smaller-sooner reward versus a larger-later reward. Whereas the delay to the small reward remains at 10 s, typically the delays to the larger reward increase in trial blocks from 10s, 20s, 40s, up to 60s. **D.** Temporal wagering tasks require animals to wait for variable delay reward following discrimination (i.e., categorization) of an uncertain stimulus. Longer wait times are generally associated with certainty, a proxy for confidence. Importantly, animals can abort the trial at any time during the delay, an outcome generally associated with uncertainty. **E.** Temporal distribution tasks require animals to discriminate between options with different distributions of reward delays. In this example, animals initiate a trial (central white square), then choose between stimuli associated with the same mean wait time (μ=10s) but different standard deviation of delays (σ either 1s or 4s).

A long-standing challenge has been to identify which parts of the brain are involved in both explicit and implicit timing, along with the neural mechanisms by which neural circuits tell time. One hypothesis is that most neural circuits have the ability to tell time if the computations those areas are involved in are time-dependent or require temporal information (Paton & Buonomano, 2018). Consistent with this hypothesis, a large number of different brain areas have been implicated in timing across a wide range of explicit and implicit tasks, including the cerebellum, striatum, parietal cortex, hippocampus, motor cortex, sensory cortex, and prefrontal cortex (Buhusi & Meck, 2005; Jennifer T. Coull, Cheng, & Meck, 2011; Issa, Tocker, Hasselmo, Heys, & Dombeck, 2020; Mauk & Buonomano, 2004; Merchant, Harrington, & Meck, 2013; Paton & Buonomano, 2018).

While it remains an open question which areas are causally responsible for timing in different tasks, there is significant evidence that different subregions of the PFC contribute to explicit timing (Bakhurin et al., 2017; Emmons et al., 2017; Kim, Ghim, Lee, & Jung, 2013; Kim, Jung, Byun, Jo, & Jung, 2009; Xu, Zhang, Dan, & Poo, 2014). For example, mPFC inactivation significantly impaired performance on a bisection task in which rats had to classify intervals as short or long (Kim et al., 2009). Additionally, large-scale extracellular recordings in OFC during a reward anticipation task, revealed a robust neural code for elapsed time from cue onset to reward (Bakhurin et al., 2017).

Overall, these findings are consistent with the notion that the PFC in general, and OFC in particular, contribute to telling time. But here we suggest that decision making requires temporal computations that extend beyond the traditional dichotomy of explicit vs. implicit timing. While standard decision making tasks often clearly include explicit and implicit timing components, decision making also requires integration of numerous time-dependent parameters. Next, we review evidence that OFC plays a crucial role in this temporal cognition.

## OFC and Temporal Discounting

Temporal discounting tasks (**Figure 1C**) typically involve offering subjects two options, one that yields a large magnitude reward and another a smaller reward (Mazur, 1987, 2007; Mazur & Biondi, 2009; Rodriguez & Logue, 1988). Generally, in animal studies, both options are initially associated with the same short reward delay, and as the task continues, the delay to the larger reward increases until the subject consistently chooses the smaller reward over the larger reward. The point at which the value of the larger-later reward is chosen equivalently to the smaller-sooner option is referred to as the indifference point in which the increased value based on reward magnitude is counterbalanced by the negative impact of increased wait time. The study of delay discounting dates back to behavioral studies in pigeons conducted in the late 1960s and early 1970s (Ainslie, 1974; Chung, 1965; Mazur, 1987; Rachlin & Green, 1972; Rodriguez & Logue, 1988). The temporal discounting rate was often modeled by adjusting the delay, but some groups also varied the amount of reward at specific delays (Mitchell, 1999; Richards, Mitchell, de Wit, & Seiden, 1997; Richards, Zhang, Mitchell, & de Wit, 1999). Temporal discounting is generally best fit by the so-called hyperbolic model of temporal discounting (K Namboodiri, Mihalas, Marton, & Hussain Shuler, 2014; Kirby, 1997; Madden, Begotka, Raiff, & Kastern, 2003; Mazur, 1987; Rachlin, Raineri, & Cross, 1991; Reynolds & Schiffbauer, 2004; Richards et al., 1997; Richards et al., 1999; Rodriguez & Logue, 1988). Specifically, the subjective value (*V*) of a reward is defined both by its magnitude (*q*) and reward delay (*d*): 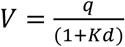, where *K* is the rate of temporal discounting. Note that according to this standard formulation, animals must 1) have an internal neural timer that allows them to measure the delay *d*; and 2) perform a computation that involves the ratio between magnitude and delay.

Since it has been established that OFC is involved in encoding the current value of reward, we further propose that OFC is embedded in a network that has access to elapsed time and can utilize this information to integrate magnitude and delay information dynamically. Indeed there is significant evidence that OFC is involved in temporal discounting in humans (Sellitto, Ciaramelli, & di Pellegrino, 2010), monkeys (Hosokawa, Kennerley, Sloan, & Wallis, 2013), and rats (Mar, Walker, Theobald, Eagle, & Robbins, 2011; Mobini et al., 2002; Peter H. Rudebeck, Walton, Smyth, Bannerman, & Rushworth, 2006). OFC lesions in both primates and rodents lead to an increase in impulsive responses in delay discounting tasks, i.e., a preference for smaller-sooner over larger-later rewards (Hosokawa et al., 2013; Kheramin et al., 2003; Mobini et al., 2002; Peter H. Rudebeck et al., 2006; Sellitto et al., 2010). However, some studies have reported that lesions of OFC lead to either no effect on choice (Abela & Chudasama, 2013; Mariano et al., 2009) or even increases in patient responses, i.e., an increase in larger-later choices (Winstanley, Theobald, Cardinal, & Robbins, 2004), with a special role for medial OFC (Mar et al., 2011).

Overall these studies suggest OFC lesions often result in more impulsive behavior. Factors contributing to this phenotype could indicate that: 1) OFC is involved in the inhibitory control necessary for delaying short-term gratification; 2) value calculations in the absence of OFC result in very strong temporal discounting because delays are overweighted; and 3) although less likely, it is possible that in the absence of OFC, the neural timers could be accelerated leading to overestimates of the actual delay—again, effectively overweighting delays. However, as mentioned above, at least two studies have reported that OFC lesions (medial OFC lesions, in particular) can shift behavior towards more patient choices. We next examine the potential methodological differences that might contribute to these discrepancies and further elucidate the role of OFC in temporal discounting and temporal cognition.

## Methodological differences in probing OFC in temporal discounting

Comparing several studies conducted on this topic in rat OFC, there appear to be no systematic differences in pre-versus post-training lesion methods that fully account for the pattern of mixed behavioral effects summarized above. Some groups administered pre-lesion training (Mar et al., 2011; Mariano et al., 2009; Peter H. Rudebeck et al., 2006; Winstanley et al., 2004), while in other studies all learning and testing phases were performed following OFC lesions (Abela & Chudasama, 2013; Kheramin et al., 2003; Mobini et al., 2002). Additionally, some groups investigated temporal discounting in mazes (Mariano et al., 2009; Peter H. Rudebeck et al., 2006), others in operant chambers with levers (Kheramin et al., 2003; Mar et al., 2011; Mobini et al., 2002; Winstanley et al., 2004) or stimuli on touchscreens (Abela & Chudasama, 2013). All but one study segregated the options spatially, instead requiring animals to associate visual cues with smaller-sooner versus larger-later rewards (Mariano et al., 2009). The results from Mariano et al. (2009) are intriguing for this reason: when rats are provided with cues during the delay period, OFC lesions do not result in an impairment in temporal discounting. This suggests that OFC may not be needed for decisions when there is a constant, salient reminder of cue-delay (or cue-outcome) associations. We discuss the importance of cues in delay-based decisions in the next section. Finally, most studies report near-identical stereotaxic coordinates for targeting whole OFC, and judging by the estimates of damage in many of these reports, there do not appear to be clear-cut differences in medial versus lateral OFC reconstructions that could explain the discordant behavioral effects. Notably, one group systematically compared medial and lateral OFC in temporal discounting and found that whole OFC lesions transiently rendered animals delay-averse (yet recovered with training), medial OFC lesions made animals delay tolerant, and lateral OFC lesions produced a delay-averse, impulsive phenotype (Mar et al., 2011), indicating there is functional heterogeneity in rat OFC for temporal discounting, as with other domains like reversal learning (Alicia Izquierdo, 2017).

In the only experiment reporting a more patient phenotype following whole OFC lesions (Winstanley et al., 2004), the lesions occurred after extensive pretraining on a temporal discounting task; the most of any study we reviewed. Additionally, a very long delay to the larger-later option was administered in this study: a maximum of 60 sec, compared to delays in the range of 10-30 sec in most other reports. And finally, this was one of the few studies where rats were (pre)trained on an integrated magnitude and delay experience within the same session. The majority of other studies (whether pre- or post-training lesions were administered) usually imposed far less training, fewer trials within each session, and trained rats to discriminate reward magnitudes *before* delays; separately not concurrently (Abela & Chudasama, 2013; Mariano et al., 2009; Peter H. Rudebeck et al., 2006). Whether animals are trained separately on magnitude versus delay (and the degree to which they have this experience with OFC online) may fundamentally change performance on temporal discounting. Indeed, Kheramin et al. (2003) derived linear functions based on choice of *d*_*A*_ (the unchanging, low magnitude reward option) versus *d*_*B*_ (the increasing delay, higher magnitude reward option), and found that their slopes were significantly steeper following OFC lesions, yet the intercept did not differ significantly between the groups. A later investigation (Kheramin et al., 2005) surmised that the parameter specifying absolute reward value was lower in OFC-lesioned animals, potentially decreasing the ability to discriminate relative reward value. One explanation for the effects observed by Winstanley et al. (2004) is that delay tolerance occurs because rats have already robustly learned to integrate delay time into the value calculation, and that after OFC lesions value estimates in downstream areas underweigh the delay because of the absence of properly integrated magnitude/delay input from OFC as it changes. OFC lesioned animals have, in effect, a deficit in updating values as the large reward delays are increased from zero to longer delays—i.e., animals essentially perseverate at the cues associated with larger rewards.

As further evidence of a role for OFC in integrating reward magnitude and delay, a modified model by Ho, Mobini, Chiang, Bradshaw, and Szabadi (1999) was used to analyze temporal discounting data following OFC lesions by (Kheramin et al., 2003). These groups proposed that the value of the reward (*V*) can be determined by the multiplicative combination of hyperbolic discounting functions for all the salient features of the reward. For the experiments described above this would be magnitude (*q*) and delay (*d*), (this was also extended to include probability or uncertainty in the original formulation): 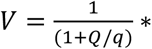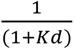, where the discounting parameters *Q* and *K* refer to the rat’s sensitivity to reward magnitude and delay, respectively. This model can explain how lesions to OFC can produce a combined effect on two discounting parameters that work in opposition, when for example there is a tradeoff between magnitude versus delay. Collectively, the data and theory point to an integration of these parameters in OFC where delay aversion or tolerance phenotypes largely depend on whether and how much animals had experience with integration of these features before OFC was taken offline. This is consistent with neural recording studies where both the encoding of time-discounted rewards and encoding of absolute reward value are found in OFC (Roesch et al., 2006).

## Neural responses in OFC to reward magnitude and time

As described earlier, several electrophysiological studies in primates point to reward (outcome) value encoding in OFC. In these studies, neuronal activity in OFC is found to be high during presentation of a particular taste-odor pairing when the animal is hungry, but is decreased once the animal is fed to satiety (Critchley & Rolls, 1996; Rolls, Sienkiewicz, & Yaxley, 1989). Neuroimaging studies in human subjects support the idea that neurons in OFC signal sensory-specific satiety (Kringelbach, O’Doherty, Rolls, & Andrews, 2003; O’Doherty et al., 2000): OFC activity only decreases when subjects are shown stimuli associated with the devalued food. Though there appears to be a dynamic value coding in OFC, is this represented at the single-cell level or the population level?

There are different explanations for how value may be represented in OFC, and we provide only a brief discussion here focused on its relevance to the representation of time. One possibility is that there is a “common scale” in OFC (a combined or multiplexed representation of value), coded as single unit activity (Montague & Berns, 2002). There is some evidence suggesting that single-cell activity does encode reward value in this way in OFC in primate (Roesch & Olson, 2005), rat (Hirokawa, Vaughan, Masset, Ott, & Kepecs, 2019; Simon, Wood, & Moghaddam, 2015) and in a pigeon functional analog of mammalian prefrontal cortex (Kalenscher et al., 2005). There is other evidence that value is stored as a population code based on electrophysiological studies in in rodents (Roesch et al., 2006; van Duuren, Lankelma, & Pennartz, 2008) and primates (Martin O’Neill & Schultz, 2010). This explanation falls in line with recent theoretical work (Buonomano & Maass, 2009; Fusi, Miller, & Rigotti, 2016) suggesting that single cell activity may appear to code certain attributes (i.e., time, magnitude), but that in effect it is the population activity as a whole coding such value. This is supported by studies that have conducted single cell recordings and then used population analyses to find that both instances code for reward value (Stott & Redish, 2014; van Duuren et al., 2009). Indeed, a recent study of both population and single unit activity in various PFC regions in nonhuman primate brain showed stimulus identity and current value codes in OFC at the population level and similar selectivity at the single-unit level (Hunt et al., 2018).

There are a handful of neural recording studies that demonstrate a correlation of elapsed time and both single cell and population activity in OFC. For example Hosokawa et al. (2013) demonstrated that single cell activity in primates undergoes larger changes in firing rate for delay-based tasks than for effort based tasks, indicating selectivity for this cost type that parallels the interference/lesion studies (Bailey, Simpson, & Balsam, 2016; Peter H. Rudebeck et al., 2006). Additionally, Roesch et al. (2006) showed that single units in rat OFC signal reward magnitude (large vs. small) and delay (short vs. long), but found no correlation between these signals. On this basis, authors concluded OFC does not integrate both magnitude and delay into a single representation (or ‘common currency’) of value, yet the task in this study did not explicitly require animals to integrate them, but rather separately presented delay and magnitude options in different blocks of trials. It should be noted that OFC may be especially important when there are discrete stimuli or cues associated with time and magnitude options, and it is perhaps in these situations where there is needed integration. In support of this, Roesch and Olson (2005) showed that activity in primate OFC *does* reflect the value of time (i.e. demonstrates integration) when cue-outcome associations are strong. Collectively, the evidence reviewed above affirm the importance of cues/stimuli in the ability of OFC to integrate magnitude and time. As reviewed above, and following results from Mariano et al. (2009), this suggests that OFC is not needed when cues are always present to signal delays or changes in delays. OFC may provide an eligibility trace to link the appearance of cues with their associated outcomes through delay periods during which cues must be represented in memory.

## OFC and Temporal Wagering

Another example of a decision making task that relies heavily on temporal cognition is the temporal wagering task (**Fig 1D**), which are often conducted in rodents, humans, and nonhuman primates (De Martino, Fleming, Garrett, & Dolan, 2013; Kepecs, Uchida, Zariwala, & Mainen, 2008; Middlebrooks & Sommer, 2012; Rolls, Grabenhorst, & Deco, 2010). In rodent temporal wagering tasks, animals are presented with perceptually ambiguous stimuli that they must categorize and report through a motor response. Following this response, a measure of confidence is estimated by how long they are willing to wait for the reward before they abort the current trial and reinitiate another. In these so-called post-decision wagering tasks, not surprisingly, animals wait longer for reward on trials with easier sensory discrimination stimuli—i.e., where they are more certain of their decision, and they are quicker to abort/reinitiate a trial when the stimuli are more uncertain. In rat studies, these stimuli can be visual (A. Stolyarova et al., 2019), olfactory (Lak et al., 2014), or auditory (Brunton, Botvinick, & Brody, 2013). In a study by Lak et al. (2014), rats received a mixture of two distinct odors at different ratios and were required to enter a nose port representing the odor that was present at a higher concentration (**Figure 1D**). Highest uncertainty in the task corresponded to trials in which odors were mixed in nearly equal proportions (48%-52%). Once rats made their response by entering the nose port, they were to stay there for a variable delay between 0.5 and 8 seconds. At any point, the rat could leave the nose port to end the trial and start a new one (indicating low confidence). In this manner, the amount of time the rat persisted in the port is a read-out of their confidence. After a high level of performance was established, investigators inactivated OFC with a GABAA agonist and found no effect on decision accuracy, but found a selective decrease in rats’ willingness to wait (measured by the normalized waiting time), i.e., the proxy of decision confidence. There is now evidence that both OFC and anterior cingulate cortex mediate decision confidence reports in rats. Interestingly, a recent study by our group (A. Stolyarova et al., 2019) found that both time wagering and reaction times to report/categorize the stimulus, though anticorrelated, reflect the certainty of the stimuli. Yet only the post-decision wagering time, not reaction time to report, is disrupted by inactivation of anterior cingulate cortex. It would be interesting to determine if OFC similarly participates in this specific way.

Temporal wagering tasks provide an interesting example of temporal cognition because it requires that animals not only maintain a running estimate of elapsed wait time, but integrate this time with the certainty of their decision. Many interesting questions arise regarding this process. For example, does integration happen right after the decision is made by setting a wait threshold that is a function of certainty, and animals abort the trial when this threshold is reached? Or rather, is there an ongoing evaluation of certainty and elapsed time, and perhaps other factors such as hunger and motivation, during the wait period? These are questions for future inquiry.

## OFC and Temporal Distributions

A final example of a temporal cognition task relates to the ability to not simply measure and represent the mean reward delays, but to encode the distribution those delays. Li and Dudman (2013) developed a novel learning task in which mice were required to approach the reward port for a water reward; the time delay between lever press and water delivery was drawn from a Gaussian probability function, and the SD selected to be narrow-to-wide (SD= 50, 750, and 2,000 ms), each with the same mean reward delay of 3 s. Interestingly, the authors found that mice use recent reward trials to infer a model of reward delay, and accumulate timing information over tens of trials to do so. In a rat study by Alexandra Stolyarova and Izquierdo (2017), subjects were given hundreds of trials of experience discriminating freely between two visual stimuli: choice of SA resulted in delay-to-reward intervals with a narrow wait time distribution (10 sec ± 1 SD) and choice of SB resulted in the same mean wait time on average, but a wider wait time distribution (10 sec ± 4 SD) (**Figure 1E)**. Aside from recording SA and SB subjective values (i.e. rats’ choices as nosepokes on the stimuli) over time, given longitudinal experience with these stimulus-delay associations, animals were also able to indicate their expectations about reward delivery, tracked experimentally by their reward port entries (i.e., the time at which, at any point in the trial, rats checked for reward). An analysis of reward port entry times revealed that rats, similar to mice, could reproduce the variance of wait times associated with individual stimuli. However, rats with ventral OFC lesions instead concentrated their reward port collection around the mean delay, indicating they lost an accurate representation of the SD, though importantly SHAM-operated rats were able to match the true distributions of those delays. The fact that rats retained a representation of the mean suggests that OFC is not needed to form simple outcome expectations based on long-term experience, but that instead it is required to accurately learn and/or represent the full temporal structure of the task over many trials (Alexandra Stolyarova & Izquierdo, 2017). Since rats received lesions of OFC prior to any training on delays, it may be of interest in future work to understand if OFC perturbation would similarly produce these changes in performance if delays were learned beforehand.

Since OFC is implicated in supporting appropriate response to changes in reward, we also sought to test how animals would respond to changes in those delay distributions (waiting less time than expected, or “upshifts” vs. waiting longer than expected, or “downshifts”). Following these manipulations, we found that experience with wider delay distributions facilitated rats’ learning for upshifts, whereas the narrower distributions facilitated rats’ learning the downshifts. This is likely accounted for by the hyperbolic shape of delay discounting, discussed above. We also surmised that positive, surprising events (i.e. near-immediate rewards when unexpected) could boost learning more strongly than negative ones. Collectively, our results provide further evidence that the representations of expected outcomes in (ventral) OFC contain information about variability in outcomes. This, we think, would allow an animal to detect changes if the events violate expectations, and indeed if these changes are meaningful and require behavioral updates (Alexandra Stolyarova & Izquierdo, 2017). These findings also fit within the broader literature implicating both ventral and lateral sectors of rat OFC in delay discounting, outcome prediction, and decision confidence (reviewed in (Alicia Izquierdo, 2017)). Learning the temporal structure of a task or temporal distributions also allows for preparation of the motor system, which manifests into faster or more consistent reaction times (MÜller-Gethmann, Ulrich, & Rinkenauer, 2003), and also allows adaptation to changing environments (Li & Dudman, 2013). Like the involvement of OFC in temporal discounting and wagering, it is less likely that OFC is involved in timing *per se* but rather the integration of timing and expected reward.

A potentially interesting parallel is the proposal that midbrain dopamine drives and enhances “anticipatory value” during reward delay periods, and areas of MFC track this value (Iigaya, Story, Kurth-Nelson, Dolan, & Dayan, 2016). Others have similarly theorized that while reward prediction errors (RPEs) are signaled clearly by dopamine transients (Schultz, 1986; Schultz, Apicella, & Ljungberg, 1993; Sharpe et al., 2017; Y. K. Takahashi et al., 2017), OFC does not itself signal RPEs (Stalnaker, Cooch, & Schoenbaum, 2015; Stalnaker, Liu, Takahashi, & Schoenbaum, 2018; Yuji K. Takahashi, Langdon, Niv, & Schoenbaum, 2016). OFC may instead be critical in constructing *expected uncertainty*, or the variance of value over repeated trials, as demonstrated by interference techniques (Soltani & Izquierdo, 2019; Alexandra Stolyarova & Izquierdo, 2017). Formally, we have defined this as the absolute value of an option, sampled over many trials (Soltani & Izquierdo, 2019). Instead of RPEs signaled on a trial-by-trial basis, OFC may be causally involved in representing the variance of time-to-reward over the entire session, or longer (i.e. across sessions). Previous versions of this argument have been put forward for OFC’s involvement in risk, or expected probabilities (M. O’Neill & Schultz, 2015).

## Conclusion

In this review, we highlight that decision making and value determination require a complex integration of information across many dimensions, and that the temporal dimension is of particular relevance because time itself is a valuable resource. Thus it is important to consider the need to measure elapsed time, generate anticipatory responses, create temporal expectations, encode previously experienced reward delays, and evaluate urgency, in decision making—we refer to the integration of these time-dependent factors as *temporal cognition*. We review evidence for the involvement of OFC in temporal cognition, and more specifically propose that the OFC contributes to the dynamic integration of reward magnitude and delay. Additionally, we note that the representation of time used to calculate value may be distinct to that of traditional timing tasks (i.e., explicit and implicit timing). The main difference is that time is not being used to simply measure a delay or generate an expectation, but rather to learn and respond to changes in the reward environment, and build a dynamic representation of value. Examples of tasks that require temporal cognition include temporal discounting, temporal wagering, and temporal distribution learning, all of which we show here depend, to some degree, on the OFC, as summarized in **Table 1**.

**Table 1.** Orbitofrontal Cortex in Temporal Cognition

| Process/Task | Description | Species | Method | Finding |
| --- | --- | --- | --- | --- |
| Explicit Timing | Discriminating between time intervals of different lengths | Humans<br>Rodents | Optogenetic inhibition, electrophysiological recordings | Inhibition of OFC impairs the ability to discriminate between two intervals. Population recordings in OFC show neural code for elapsed time |
| Implicit Timing* | Learning the temporal structure of a task improves subsequent performance | Humans | Psychophysics | Enhanced performance when the stimulus-target interval matches the expected interval |
| Temporal Discounting | Choosing between a smaller, sooner vs. larger, later reward | Humans<br>Monkeys<br>Rodents | Lesions, transient inactivations, and electrophysiological recordings | OFC lesions mostly result in delay-aversion; mixed results depend on experience integrating reward magnitude and time, and presence of cues |
| Temporal Wagering | Post-decision waiting for reward vs. aborting/reinitiating the trial | Humans<br>Monkeys<br>Rodents | Lesions, transient inactivations, and electrophysiological recordings | OFC inactivation impairs willingness-to-wait (decision confidence) without affecting decision accuracy |
| Temporal Distributions | Discriminating different delay distributions | Rodents | Lesions | OFC lesions produce a decreased ability to accurately represent variability in reward outcomes |

\*No known experiments specifically testing the role of OFC in implicit timing.

We emphasize the need for future studies that systematically manipulate the level of integration of magnitude and delay and probe OFC involvement. For example, if delays are discriminated separately from magnitude, is OFC similarly important as when delays and magnitude are learned in combination? Additionally, what systems support this multiplexing of magnitude and delay for a ‘common currency’ in OFC? Amygdala and striatum are excellent candidates, yet circuit mapping is needed to determine this. Another area of research that may prove worthwhile would be to determine the functional heterogeneity of temporal cognition in OFC, as several groups have discovered rich differences in medial versus lateral OFC in other forms of cognition.

## Acknowledgements

We thank Madeline Valdez for her help with the literature search.

## Notes

**Funding** This work was supported by R01 DA047870 (Izquierdo) and UCLA Division of Life Sciences Retention Funds (Izquierdo).

**Disclosure** There is no conflict of interest.

### Competing Interest Statement

The authors have declared no competing interest.

